# Red light responsive Cre recombinase for bacterial optogenetics

**DOI:** 10.1101/2024.05.30.596707

**Authors:** Fereshteh Jafarbeglou, Mary J. Dunlop

## Abstract

Optogenetic tools have been used in a wide range of microbial engineering applications that benefit from the tunable, spatiotemporal control that light affords. However, the majority of current optogenetic constructs for bacteria respond to blue light, limiting the potential for multichromatic control. In addition, other wavelengths offer potential benefits over blue light, including improved penetration of dense cultures and reduced potential for toxicity. In this study, we introduce OptoCre-REDMAP, a red light inducible Cre recombinase system in *Escherichia coli*. This system harnesses the plant photoreceptors PhyA and FHY1 and a split version of Cre recombinase to achieve precise control over gene expression and DNA excision. We optimized the design by modifying the start codon of Cre and characterized the impact of different levels of induction to find conditions that produced minimal basal expression in the dark and induced full activation within four hours of red light exposure. We characterized the system’s sensitivity to ambient light, red light intensity, and exposure time, finding OptoCre-REDMAP to be reliable and flexible across a range of conditions. In co-culture experiments with OptoCre-REDMAP and the blue light responsive OptoCre-VVD, we found that the systems responded orthogonally to red and blue light inputs. Direct comparisons between red and blue light induction with OptoCre-REDMAP and OptoCre-VVD demonstrated the superior penetration properties of red light. OptoCre-REDMAP’s robust and selective response to red light makes it suitable for advanced synthetic biology applications, particularly those requiring precise multichromatic control.

## Introduction

Optogenetic approaches can be used to manipulate cellular processes for light-mediated control, offering tunable and precise modulation of gene expression^1–10^. Unlike traditional chemical inducers, light is straightforward to program, it can be added and removed rapidly, and it can be spatially patterned, providing the potential for spatiotemporal control and dynamic perturbations without the need for media alterations or system disruptions. Light is also orthogonal to many endogenous cellular processes, allowing optogenetic tools to control cellular functions with minimal cross-talk or off-target effects^11^. Moreover, because light is easily programmed, optogenetic tools can also interface with dynamic computer-based control and feedback^12–16^. These benefits have led to the use of cellular optogenetic tools in a wide range of applications, from basic research elucidating fundamental biological mechanisms^17–19^ to advanced biotechnological and therapeutic interventions^20,21^.

Recent efforts in microbial optogenetics have capitalized on the precise and programmable nature of light. For example, light-sensitive proteins have been used to modulate enzyme expression for biofuel production^22,23^, to coordinate bacterial growth via metabolic regulation^24,25^, and to inhibit the formation of biofilms^26^. In addition, optogenetic techniques have enabled light-induced drug release from hydrogels^27^, precise positioning of *E. coli* on substrates^28^, and the production of biomaterials^29–31^.

Optogenetic systems exist that respond to wavelengths ranging from near-infrared to ultraviolet light^32–35^. The majority of existing bacterial optogenetic systems are blue light responsive, and one advantage of this is that the chromophores required for their light-mediated response are flavin nucleotides, which are endogenous to all bacteria^11^. However, blue light is more prone to phototoxicity than other wavelengths, introducing the potential for cellular damage^36^. It is also more susceptible to scattering and absorption by biological molecules, leading to less precise spatial targeting of specific cell populations^36^. We chose to focus here on developing a red light responsive system. Red light has superior penetration capabilities compared to blue light, allowing for deeper stimulation within biological tissues^37^ or within bioreactors. Moreover, having both blue and red light inducible systems could allow the design of circuits with orthogonal light inputs for multichromatic control.

In plants, the capacity to perceive and respond to red/far-red light relies on proteins from the phytochrome photoreceptor family^38^. For example, PhyA is a photoreceptor from *Arabidopsis thaliana*^38^ that binds covalently to the chromophore phycocyanobilin (PCB)^39^, resulting in a photochemically functional photoreceptor that is responsive to red light. In the native plant context, after receiving red light PhyA is translocated into the nucleus through interactions with shuttle proteins, including FHY1^40^. Upon red light illumination (λ_max_ = 660 nm), PhyA and FHY1 associate, whereas under far-red light (λ_max_ = 730 nm), they dissociate. Taking advantage of this red light inducible protein interaction, Zhou et al.^41^ developed and optimized a compact and sensitive red light mediated photo switch, denoted ‘REDMAP’, for control of transcription in mammalian cells. Using this design, they fused PhyA to the yeast Gal4 DNA-binding domain and FHY1 to a tetrameric repeat of the minimal herpes simplex virus-derived transcription activator (VP16) to form a light-responsive transcription factor. They demonstrated that after red light induction, PhyA and FHY1 associate, resulting in the formation of a functional transcription factor complex (Gal4-VP16). They tested different REDMAP system compositions and observed that a truncated version of PhyA comprised of amino acids 1–617 (which they denote ‘ΔPhyA’), resulted in a 13-fold increase in red light induced gene expression via Gal4–VP16^41^.

In this study, we sought to use light to control recombinase activity. Recombinases are enzymes capable of targeting specific 30−50 base pair sequences within DNA and modifying the designated DNA flanked by these recognition sites. For example, Cre is a widely used tyrosine recombinase that is derived from the P1 bacteriophage. Cre can recognize DNA flanked by loxP sites and, depending on the orientation of the loxP sites, will excise, invert, or translocate the DNA sequence^42^. The proficiency of recombinases in DNA manipulation has been widely recognized for synthetic biology applications, where they have been used to construct complex cellular logic circuits^43^ and engineer gene circuits with memory^44,45^. Researchers have also recognized the potential of combining optogenetic control with recombinase function. For example, light-responsive recombinases have been used in mammalian systems^43,46–49^ and yeast^50^. Optogenetic recombinases have also been developed for bacteria^3,51,52^, where they have been used to regulate the expression of antibiotic resistance genes^51,53^ and to directly capture spatial information and input signals via light using DNA as a means to store digital information^54^. Inducible recombinase designs typically employ a split protein strategy, where the recombinase is rendered light-sensitive by splitting the gene into N-terminal and C-terminal fragments, each linked to a light-responsive photodimer or photoreceptor sequence^2,3,43^. Upon light stimulation, conformational changes in the photodimer or association of photoreceptor proteins promote the reassembly of the two recombinase fragments to form a functional enzyme.

In this work, we developed and optimized a red light inducible Cre recombinase in *E. coli*. We began our efforts with OptoCre-VVD, a blue light inducible Cre variant that we developed and optimized in a previous project^3^. By integrating red light responsive FHY1 and ΔPhyA photoreceptors^49^ into our design to replace the blue light responsive VVD photodimers, we engineered a novel red light responsive Cre. Here, we introduce this design, which we call ‘OptoCre-REDMAP’. First, we investigated the robustness of the system in response to different split Cre induction levels. Our initial design exhibited undesirable basal activity in the dark, which we reduced by optimizing the Cre start codon. The new construct showed robust activation with negligible basal expression. Then, we characterized sensitivity of the OptoCre-REDMAP construct to ambient light exposure, variations in red light intensity, and light exposure time. To provide examples of future applications, we used this system to activate an antibiotic resistance gene in cells and used co-culture experiments to test OptoCre-REDMAP alongside the blue light inducible OptoCre-VVD system to investigate its potential for multichromatic control. Additionally, we demonstrated that red light allowed for effective induction of OptoCre-REDMAP even when cultures were partially obscured, a condition under which blue light responsive OptoCre-VVD showed markedly reduced activation. Overall, OptoCre-REDMAP is a robust red light responsive system, which is suitable for multichromatic control in bacterial synthetic biology applications.

## Results

To render Cre recombinase red light activatable, we started with the blue light inducible OptoCre-VVD^3^, which is comprised of N-terminal and C-terminal fragments of Cre fused with VVD photodimers. Blue light induces a conformational change in VVD to allow dimerization, bringing the Cre fragments together^3^. We constructed OptoCre-REDMAP by replacing the VVD photodimers with the FHY1 and ΔPhyA photoreceptors, attaching FHY1 to the N-terminal fragment of the Cre sequence, and ΔPhyA to the C-terminal fragment (Fig. 1). We placed the two components of OptoCre-REDMAP together on an operon, with a ribosome binding site separating nCre-FHY1 and cCre-ΔPhyA. We employed an IPTG-inducible P_lacUV5_ promoter to regulate the expression of split Cre to prevent premature target DNA excision prior to IPTG induction (Fig. 1). Additionally, ΔPhyA requires the PCB chromophore for photochemical activity, thus we included the *ho1-pcyA* gene that produces PCB in the circuit design (Methods).

**Figure 1.**
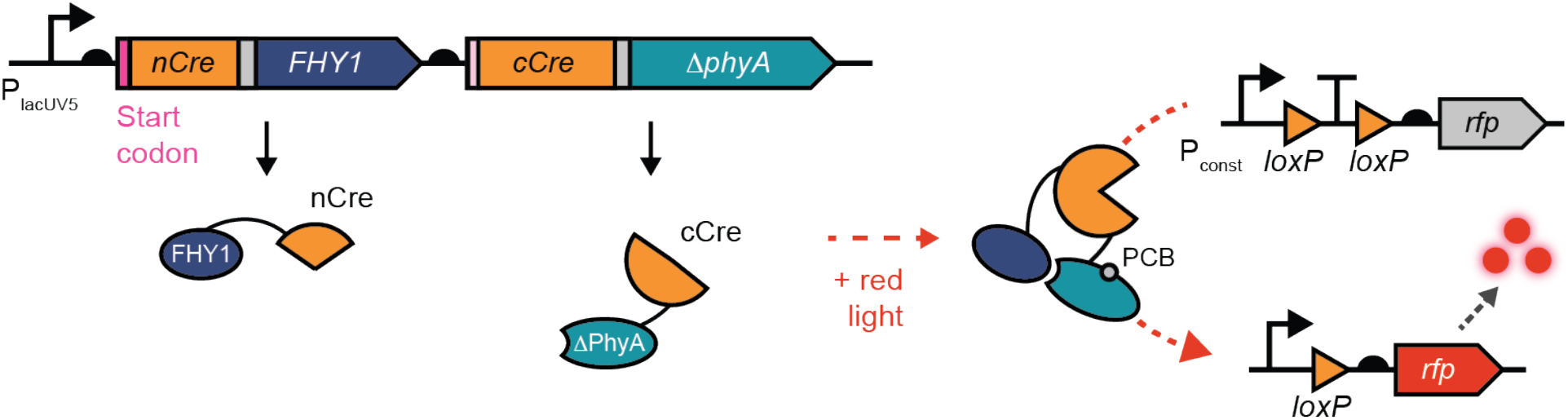
Red light induced recombination in *E. coli*. Split Cre fragments are linked to FHY1 and ΔPhyA photoreceptors and expressed under the control of an IPTG-inducible promoter (P_lacUV5_). ΔPhyA must be covalently bound to the chromophore phycocyanobilin (PCB) to be photochemically active. When exposed to red light, FHY1 and ΔPhyA associate, forming a functional Cre protein. Cre can then act on the reporter plasmid, excising the loxP-flanked transcriptional terminator and allowing expression of RFP under the control of a constitutive promoter (P_const_).

Cre recognizes and excises DNA flanked by loxP sites that are oriented in the same direction. To test the function of OptoCre-REDMAP, we used a reporter where a transcriptional terminator flanked by loxP sites is placed between a constitutive promoter and the gene encoding red fluorescent protein (RFP). In the absence of functional Cre, the terminator effectively halts the transcription process and prevents the expression of RFP. With functional Cre, the terminator is excised, allowing transcription and subsequent production of RFP (Fig. 1).

To test the performance of OptoCre-REDMAP, we exposed cultures to red light for 4 hours. We kept another set of samples in the dark during the same period (Fig. 2a). For comparison, we also included a negative control strain that only contained the reporter plasmid and a positive control strain consisting of the reporter plasmid with the transcriptional terminator excised, allowing full expression of RFP. We found that OptoCre-REDMAP was functional, with clear differences in fluorescence between the dark and red light conditions (Fig. 2a). We conducted experiments using a range of IPTG concentrations, finding that full activation was achieved for IPTG concentrations at or above 50 μM. However, our construct exhibited significantly higher basal expression than the negative control for most levels of IPTG induction, indicative of leaky OFF-state behavior in the dark. First, we attempted to decrease the leaky activity of the system by reducing the duration of light induction to 2 hours. This led to a modest decrease in the leaky activity, in addition to a slight decrease in the red light induced levels of RFP (Fig. 2b). However, we ultimately deemed this approach insufficient given the residual leaky activity in the dark conditions. We also tried to further reduce the red light exposure time, but with shorter red light induction the system did not reach full activation (Fig. S1a). Next, we tried to decrease the red light intensity rather than its duration. However, we found that the red light intensity did not significantly impact the behavior of the system and thus was not a viable strategy for decreasing the basal expression (Fig. S1b).

**Figure 2.**
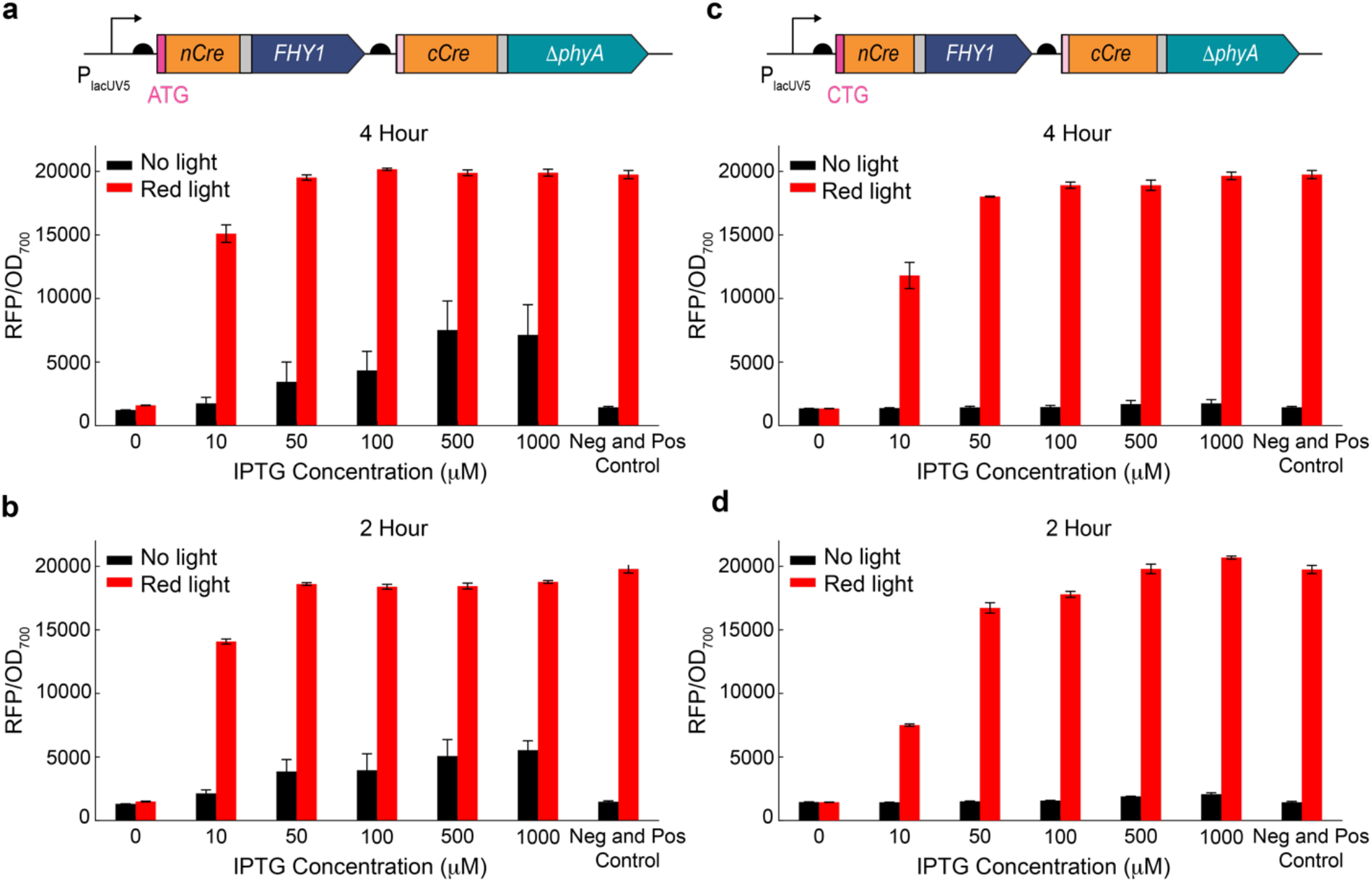
Comparison of two different designs for OptoCre-REDMAP. (a) RFP expression levels of OptoCre-REDMAP (ATG version) at different IPTG concentrations with 4 hours and (b) 2 hours of red light exposure. (c) RFP expression levels of OptoCre-REDMAP (CTG version) at different IPTG concentrations with 4 hours and (d) 2 hours of red light exposure. In all cases, error bars show standard deviation around the mean (n=3 biological replicates).

Because straightforward changes based on IPTG or red light induction were insufficient to reduce basal expression while maintaining light induced activity of OptoCre-REDMAP, we next considered whether aspects of our construct design could be changed to improve performance. First, we tried two alternative split sites, located at amino acid 136 or 270 in the Cre sequence. We selected these locations based on their fold change in response to light in related studies on optogenetic split protein designs by Weinberger et al.^43^ and Tague et al.^52^. However, both designs performed poorly relative to the original split site at amino acid 43 (Fig. S1c). Working with the original split location, we next focused on reducing basal expression of OptoCre-REDMAP by changing the start codon of nCre-FHY1 from ATG to CTG. Hecht et al.^55^ measured the translation initiation rate corresponding to 64 different start codons in *E. coli* and observed that CTG exhibits 100 times lower expression than ATG. Therefore, we hypothesized that changing the start codon from ATG to CTG would help reduce RFP expression in the dark. By switching the start codon, the total number of split Cre fragments in the cell will decrease, ideally resulting in conditions where the number of split Cre proteins capable of association will be insufficient to activate in the dark, yet adequate for achieving full activation when fragments combine in the presence of red light. We constructed the new variant and quantified RFP expression at different IPTG concentrations. The new design exhibited substantially reduced basal expression relative to the ATG version, while maintaining high RFP levels when exposed to red light (Fig. 2c). The basal expression was minimal for all IPTG concentrations we tested, remaining at levels comparable to the negative control. We found that full activation by OptoCre-REDMAP can be achieved with IPTG concentrations at or above 50 μM with 4 hours of red light illumination. However, if a shorter red light illumination period, i.e. 2 hours, is desirable, increasing the IPTG concentration to 500 μM will result in full activation (Fig. 2d). Given the minimal basal expression and robust activation under red light, we proceeded with this optimized construct for subsequent experiments.

We tested the impact of red light intensity and exposure time on the performance of the CTG variant of OptoCre-REDMAP. First, we confirmed that light induction did not impact growth. When conducting experiments to optimize the behavior of OptoCre-REDMAP in Fig. 2, we used red light with full intensity (155 μW/cm^2^) to activate the system. By analyzing both the optical density values (Fig. 3a) and microscopy images of individual cells (Fig. S2a-b) at 155 μW⁄cm^2^, we observed no notable disparities in growth rates or cell morphology between experimental conditions with and without light exposure, indicative of minimal phototoxicity effects with this exposure level. Next, we assessed whether results were sensitive to light induction levels by testing a range of intensities, spanning 15.5 μW/cm^2^, a very dim lighting condition, to 155 μW/cm^2^, bright lighting (Fig. 3b-c). We found that the light intensity in this range did not significantly affect the behavior of our system with 4 hours of light exposure (Fig. 3b). With 2 hours of light, we observed modest decreases in fluorescence levels for the low light intensity values (Fig. 3c), but overall, the system responds robustly to many levels of red light. We also tested very dim light conditions, measuring responses to levels below 15.5 μW/cm^2^ and observed graded activation in this regime (Fig. S2c).

**Figure 3.**
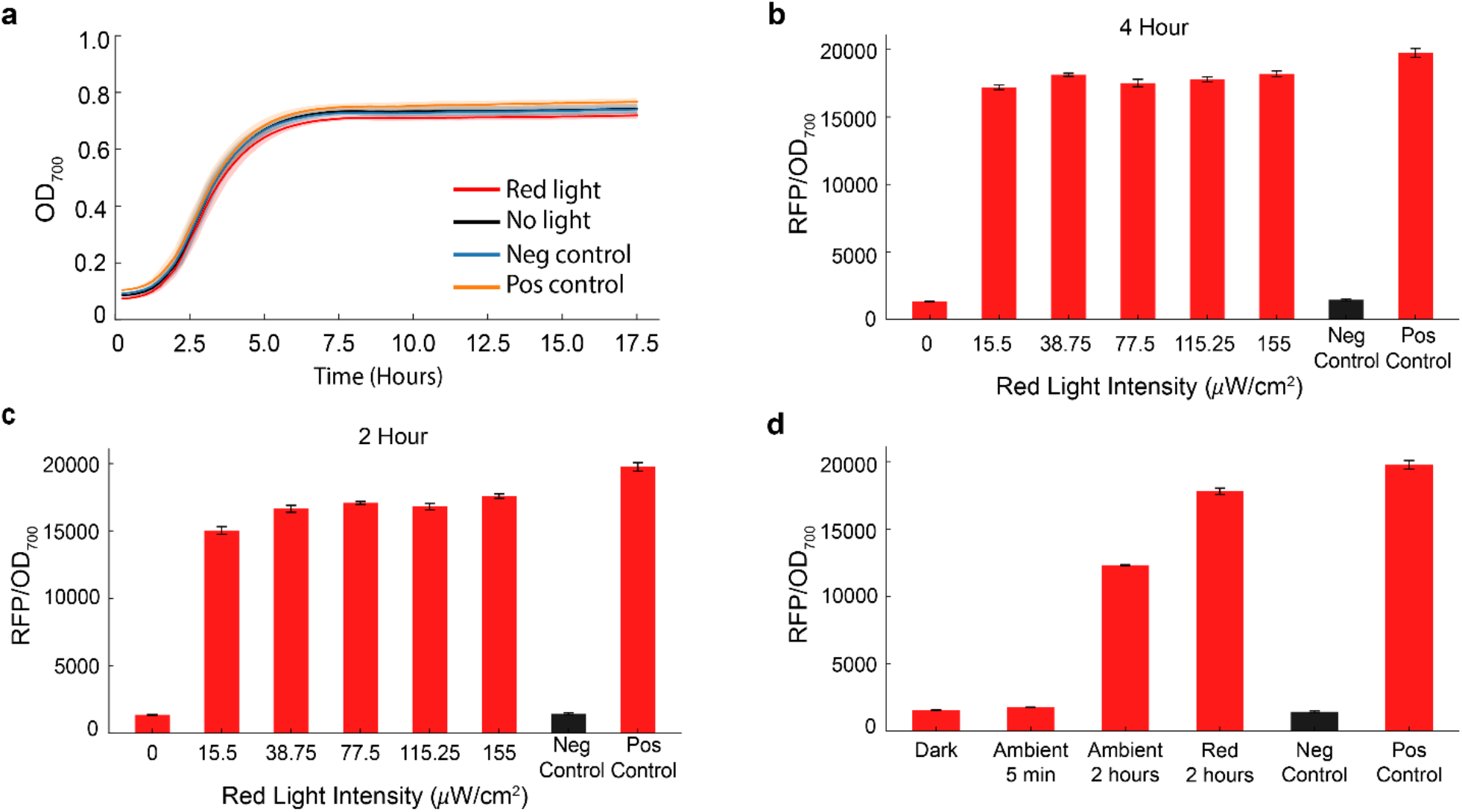
Characterizing the behavior of OptoCre-REDMAP. (a) Optical density at 700 nm, OD700, values for cultures subjected to 4 hours of red light compared to the dark state, negative, and positive control. Shaded error bars represent standard deviation around the mean from plate reader data (n=3 biological replicates). (b) RFP output in response to red light intensities ranging from 15.5 to 155 μW⁄cm^2^ with 4 hours or (c) 2 hours of light exposure. (d) RFP expression of OptoCre-REDMAP in response to 5 minutes and 2 hours of exposure to ambient light compared to the dark state, 2 hours of red light exposure, and the negative and positive control. Error bars show standard deviation around the mean (n=3 biological replicates).

Next, we conducted tests to characterize the sensitivity of OptoCre-REDMAP to ambient light exposure. When working with light-responsive systems, it is important to assess their sensitivity to surrounding light conditions since short-term exposure to ambient light is inevitable in practical experimental conditions where plates are temporarily removed from the dark, as might occur during the transfer of cultures for flow cytometry or microscopy assays. In addition, it would be helpful to know whether keeping the cultures in the dark is necessary for the robustness of the results. Measuring light conditions in the laboratory, we found that ambient intensities were ∼3.6 μW/cm^2^ at a wavelength of 630 nm. We assessed the performance of OptoCre-REDMAP under a brief, 5-minute exposure to ambient light, and a long, 2-hour exposure to ambient light (Fig. 3d). We found that OptoCre-REDMAP is not very sensitive to short periods of ambient light, suggesting its reliability in conditions with incidental exposure (Fig. 3d). However, the 2-hour ambient light exposure produced substantial activation of the system, indicating that keeping the cultures in the dark for most portions of the experiment is necessary for reproducibility and robustness of the results (Fig. 3d).

In addition to assessing the ON and OFF states of our system after red light stimulation, we aimed to investigate the real-time dynamics of activation in light-inducible cells relative to both negative and positive controls. The purpose of this experiment was to monitor fluorescence levels continuously without the delay introduced by overnight incubation in the dark, which is typically used to allow protein expression and maturation. To achieve this, we diluted the cultures 1:100 and transferred them to a plate reader immediately following light stimulation. We tracked RFP expression over an 18-hour period and found that RFP expression in the light-activated cells rapidly increased to levels comparable to the positive control, indicating that RFP was fully induced at the conclusion of the 4 hour light stimulation period (Fig. S3).

Next, we sought to examine the impact of varying durations of red light exposure on cells by characterizing DNA excision and quantifying RFP expression levels in both bulk culture and at the single cell level. To investigate this, we systematically varied the duration of red light stimulation, ranging from 5 minutes to 8 hours, and assessed activation at both the bulk culture and single-cell levels. We first evaluated the extent of DNA excision at the bulk culture level using PCR and gel electrophoresis. We observed a clear shift in band patterns indicating successful excision of the transcriptional terminator after a minimum of 2 hours of red light exposure (Fig. 4a), consistent with our fluorescence measurements. These results confirmed that 2 hours of red light was sufficient to activate the majority of cells within the population. To quantify RFP expression levels, we measured fluorescence in bulk culture samples (Fig. 4b). The results indicated that cells exposed to red light for 2 hours or longer achieved RFP expression levels comparable to those of the positive control, suggesting full activation of the system. We further extended our analysis to the single-cell level to evaluate whether these bulk culture trends were representative of individual cell behavior using single cell microscopy. After bulk culture light stimulation and overnight incubation, we placed cells on agar pads and observed cells individually (Fig. 4c-d). Our observations revealed that while there was some variation in RFP expression intensity between individual cells (Fig. S4), the overall activation patterns aligned closely with the bulk culture data. Specifically, the majority of single cells exhibited RFP expression following a 2 hour or longer light exposure, validating that this duration was sufficient to achieve high levels of activation across the population.

**Figure 4.**
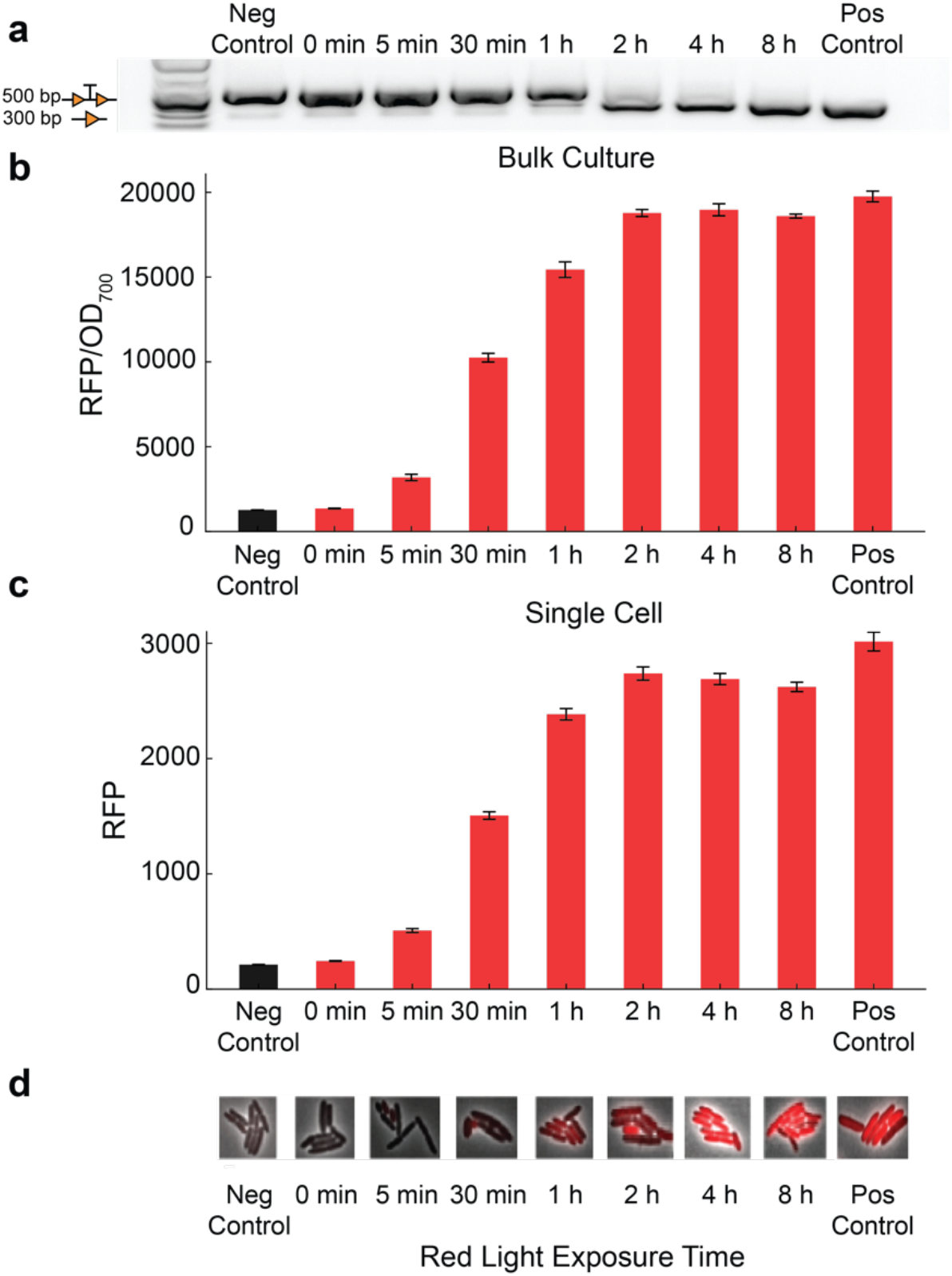
Light exposure duration necessary to induce excision by OptoCre-REDMAP and comparison of bulk culture and single cell results. (a) DNA gel image showing reporter bands with and without transcription terminator excision. (b) Bulk culture fluorescence measurement of RFP after red light exposure with different durations. For illumination durations shorter than 8 hours, the cultures were kept in the dark for the remainder of the 8 hours. The error bars show standard deviation around mean value (n=3 biological replicates). (c) Single cell fluorescence microscopy averages of RFP values. Error bars show standard error around the mean value (n ≈ 800 single cells) (d) representative microscopy images (scale bar = 2 μm).

To demonstrate the versatility of our system in activating genes beyond fluorescent reporters, we substituted *rfp* for a gene encoding chloramphenicol acetyltransferase (*cat*), which provides enzymatic resistance against chloramphenicol^56^. Following this modification, we conducted a minimum inhibitory concentration (MIC) assay to assess whether the OptoCre-REDMAP system could successfully activate *cat* in response to red light stimulation, resulting in antibiotic resistant cells. Cells exposed to red light demonstrated significantly higher resistance to chloramphenicol compared to the negative control (Fig. 5), indicating that the *cat* gene was efficiently expressed. Importantly, cells that were kept in the dark showed resistance levels similar to the negative control, confirming that the system has minimal leaky expression. This feature makes it particularly suitable for MIC assays and other applications that require stringent control over expression.

**Figure 5.**
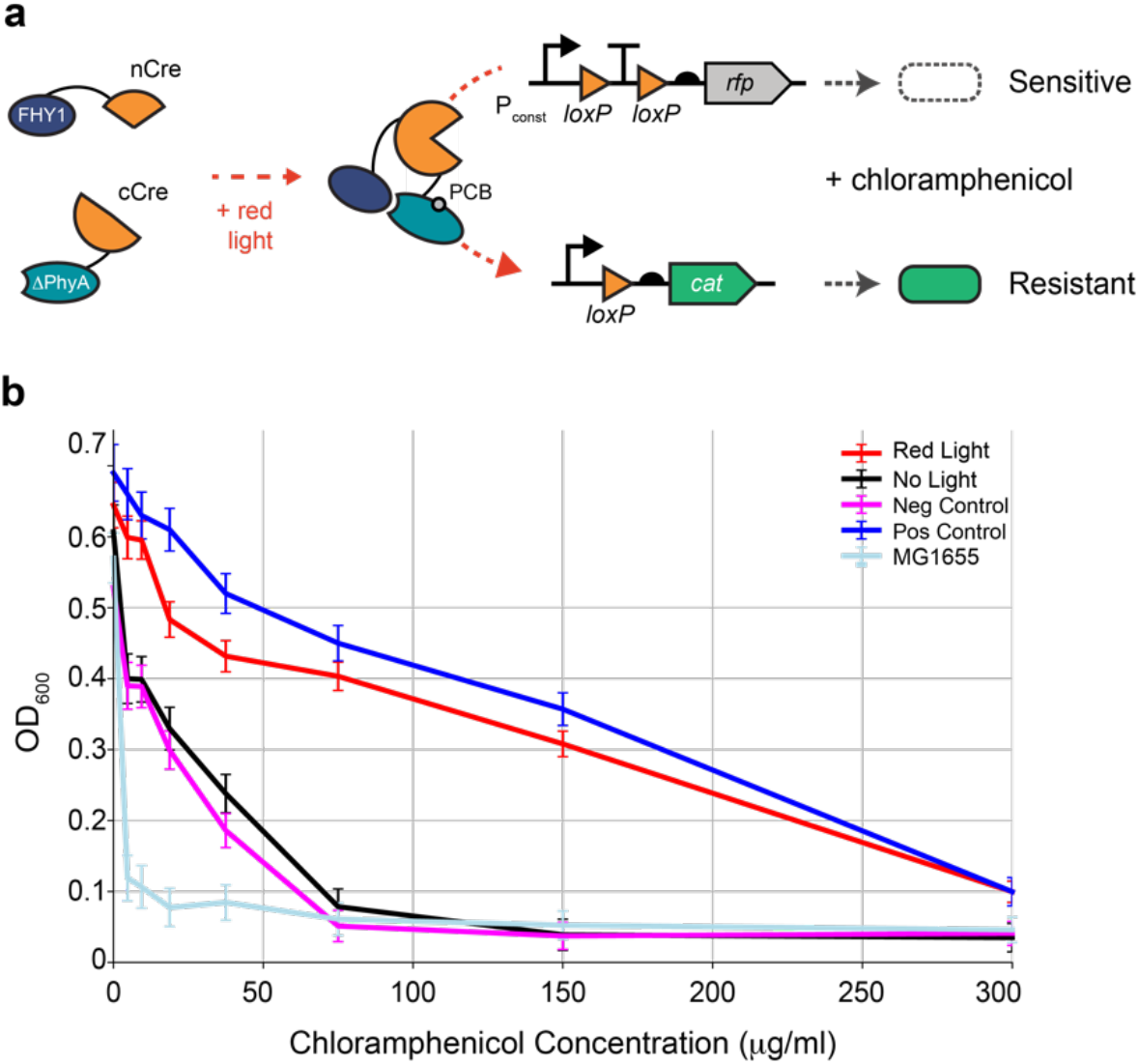
Minimum inhibitory concentration (MIC) curves of the cells grown in chloramphenicol overnight. (a) Schematic of the OptoCre-REDMAP system. When exposed to red light Cre becomes active and can excise a transcription terminator between two loxP domains, allowing increased expression of chloramphenicol acetyltransferase (*cat*). (b) Growth measured by OD600. The strains in both the dark and light conditions contain the *cat* construct and OptoCre-REDMAP. Negative control cells contain the *cat* construct, but no Cre recombinase. Positive control cells contain a constitutively expressed *cat* construct with transcriptional terminator excised. MG1655 cells contain no constructs. The error bars represent standard deviation around the mean from plate reader data (n = 3 biological replicates).

To support future applications employing multichromatic control, we characterized the performance of red light activated OptoCre-REDMAP together with blue light activated OptoCre-VVD. We used OptoCre-REDMAP to activate expression of RFP and OptoCre-VVD to activate GFP and subjected cells to red and blue light inputs. First, we conducted experiments where we grew the cell cultures separately and tested for induction in the presence of red or blue light. To achieve this, we subjected each system to red or blue light stimulation for 4 hours across a range of IPTG concentrations. Our aim was to determine the optimal experimental conditions conducive to activating both constructs under their respective light stimuli. Both constructs remained consistently inactive across all IPTG concentrations when subjected to opposite light stimulation, with OptoCre-REDMAP responding to red light and not blue and OptoCre-VVD to blue but not red (Fig. 6a-b). Next, we created co-cultures where we mixed cells with OptoCre-REDMAP and OptoCre-VVD together to test our ability to selectively activate expression. Consistent with the single culture findings, OptoCre-REDMAP responded exclusively to red light and remained inactive under blue light, while OptoCre-VVD was selectively activated by blue light and not red in the co-culture experiment (Fig. 6c). Overall fluorescence levels were lower in the co-culture experiments than in the single culture experiments, as expected due to the presence of the other strain. In addition, we found that full induction required higher concentrations of IPTG than in the single culture case for both strains. These single and co-culture results demonstrate the potential for orthogonal multichromatic control of distinct genetic programs in mixed populations, without interference between the systems.

**Figure 6.**
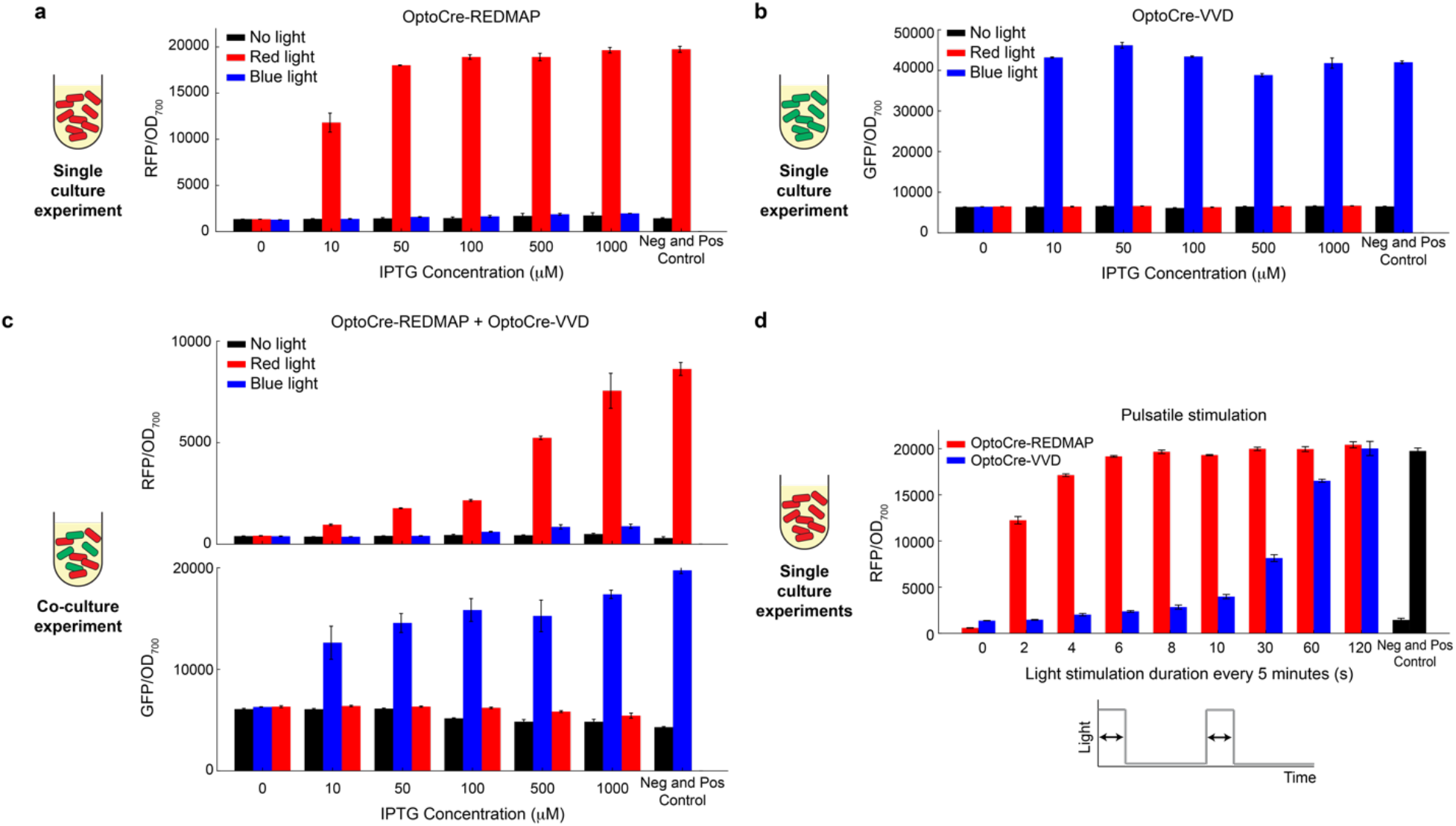
OptoCre-REDMAP and OptoCre-VVD respond orthogonally to red and blue light inputs in single cultures and co-cultures. (a) RFP levels of OptoCre-REDMAP subjected to 4 hours of constant dark, red light, or blue light. (b) GFP levels of OptoCre-VVD subjected to 4 hours of constant dark, red light, or blue light. (c) Co-culture experiment showing RFP and GFP levels of the OptoCre-REDMAP and OptoCre-VVD co-culture subjected to 4 hours of constant dark, red light, or blue light. (d) RFP levels of OptoCre-REDMAP and OptoCre-VVD single cultures subjected to pulses of red or blue light with varying durations. To make the fluorescence levels comparable to each other, we used the same RFP reporter plasmid construct for both OptoCre-REDMAP and OptoCre-VVD. In all cases, the negative control is a strain with only the reporter plasmid with the terminator unexcised and the positive control is a strain with only the reporter plasmid where the terminator is fully excised. In all cases, the error bars show standard deviation about the mean (n=3 biological replicates).

We were also interested in determining the shortest light stimulation pulse required to activate the systems, which is useful in setups where prolonged constant light stimulation is not feasible and the systems can only be stimulated in a pulse-like manner. For example, in time-lapse microscopy, multiple fields of view may need to be illuminated sequentially, and optogenetic stimulation must also be OFF during intervals when fluorescent protein levels are measured. To investigate the light stimulation requirements, we exposed the OptoCre-REDMAP and OptoCre-VVD systems to pulses of red or blue light of varying durations (e.g., 0-120s) delivered every 5 minutes over a total period of 18 hours in single culture experiments. We found that a 6-second pulse of red light every 5 minutes was sufficient to activate OptoCre-REDMAP, while OptoCre-VVD required a 60-second pulse of blue light for reliable activation (Fig. 6d). These findings help refine the parameters for optimal use of the systems in applications where light stimulation must be limited.

Finally, to demonstrate the advantage of using red light for deeper penetration, such as what would be required within biological tissues or dense cultures within bioreactors, we tested the performance of the OptoCre-REDMAP and OptoCre-VVD systems using a skin-mimicking phantom. The phantom, which was designed to mimic the optical absorption and scattering properties of human skin (Table S1), was used to assess how light penetration affects system activation. We utilized a thin 1mm phantom and placed it between the red or blue light LEDs and the well plates containing the cultures (Fig. 7a). We found that red light exhibited significantly better penetration compared to blue light, allowing for robust activation of OptoCre-REDMAP even through the phantom (Fig. 7b). In contrast, blue light’s higher absorption and scattering properties resulted in a marked reduction in activation efficiency for OptoCre-VVD (Fig. 7c). These results highlight one of the major advantages of using red light in optogenetic systems, particularly for applications requiring activation in denser cultures or at greater depths, where red light’s superior penetration is critical.

**Figure 7.**
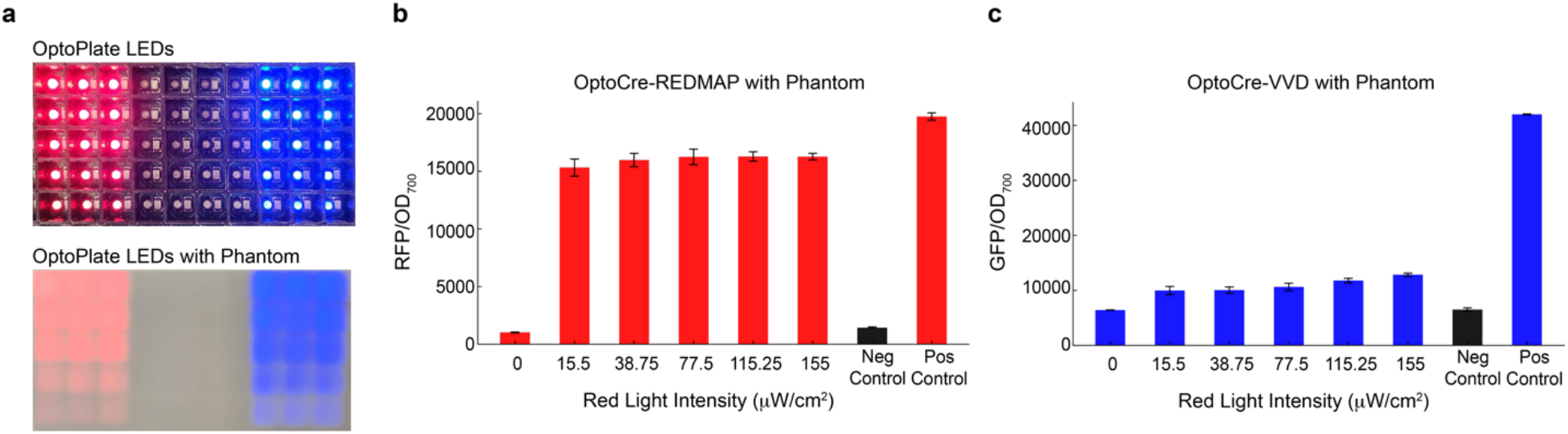
Light penetration tests with skin-mimicking phantom partially obscuring light from LEDs. (a) Image of the OptoPlate-96 wells with red and blue LEDs at different intensities and the same device covered with a phantom. (b) RFP output of OptoCre-REDMAP in response to red light intensities ranging from 15.5 to 155 μW⁄cm^2^ with 4 hours of light exposure when using the phantom. (c) GFP output of OptoCre-VVD in response blue light intensities ranging from 15.5 to 155 μW⁄cm^2^ with 4 hours of light exposure when using the phantom. In all cases, the error bars show standard deviation about the mean (n=3 biological replicates).

## Discussion

In this study, we developed, optimized, and characterized a red light activatable Cre recombinase for *E. coli*. Leveraging the FHY1-ΔPhyA light mediated protein interaction^41^, we engineered Cre recombinase to respond to red light stimuli. After testing the resulting construct, OptoCre-REDMAP, at various IPTG concentrations we observed that while photochemically active, the construct showed high basal expression, limiting its robustness. Following attempts to mitigate basal expression by adjusting exposure time and light intensity, and testing two alternative protein split sites, we ultimately modified the genetic circuit design by changing the start codon of nCre-FHY1 from ATG to CTG. This modification led to a substantial reduction in basal expression while preserving red light induced activity. Importantly, our new construct demonstrated robust and reproducible performance across varying light durations, IPTG concentrations, and red light intensities. Notably, red light exposures of 2 to 4 hours with IPTG induction were sufficient to achieve full activation. To further expand the utility of OptoCre-REDMAP, we conducted co-culture experiments to explore its potential in multichromatic control alongside the blue light responsive OptoCre-VVD system. In these experiments, we showed that OptoCre-REDMAP is selectively activated by red light without cross-activation by blue light, while OptoCre-VVD responds only to blue light, demonstrating orthogonal control in a mixed population. This co-culture setup highlights the feasibility of multiplexing gene regulatory networks in bacterial systems using distinct light inputs for independent control of different genetic programs. Additionally, we demonstrated how red light’s superior penetration properties allowed for light induction of cultures even through a skin-mimicking phantom that partially obscured the light, while induction of OptoCre-VVD with blue light was severely curtailed. This characteristic makes OptoCre-REDMAP particularly suitable for applications involving dense cultures or complex biological environments, where light penetration is crucial.

While OptoCre-REDMAP shows great promise, there are areas for further research and improvement. For instance, exploring different recombinase variants, such as Flp may enhance the system’s performance and expand its applications^43^. Alternative recombinases could also allow two constructs that are controlled by different light signals to be used in the same cell, provided the recognition sites are orthogonal. In addition, it would be interesting to test how light wavelengths for other bacterial optogenetic designs that do not use blue light as an input, such as CcaS-CcaR^34,57^ and UirS−UirR^32^, impact OptoCre-REDMAP. Further, we have demonstrated that activation kinetics differ between OptoCre-REDMAP and OptoCre-VVD and engineering variants that change the response times could help to alter the sensitivity of the new designs.

In conclusion, OptoCre-REDMAP offers robust and reliable activation across a range of red light intensities and exposure times while remaining insensitive to blue light, making it amenable to applications that require multichromatic control. In addition, its capacity for deeper penetration compared to blue light-based systems makes it a promising tool for future optogenetic studies in applications involving biological tissues or dense cultures.

## Methods

### Strains and Plasmids

In this study, we used *E. coli* strain MG1655. All recombinase constructs use a plasmid with a low copy pSC101 origin of replication and an ampicillin resistance gene. The recombinase genes are under the control of the IPTG-inducible P_lacUV5_ promoter (plasmid map in Fig. S5a). The construct was derived from the pBbS5a BglBrick plasmid^58^. All reporter plasmids use a medium copy p15A origin of replication with a kanamycin resistance gene and the gene for red fluorescent protein (mCherry) under the control of a constitutive, medium strength promoter (P_W4_). The P_W4_ promoter is modified from the phage T7 A1 promoter (ATCGTTTAGGCACCCCAGGCTTTACACTTTATGCTTCCGGCTCGTATAATGTGTGGAATTGTGAGCGGATAACAAT)^59^. The PCB encoding gene, *ho1-pcy*, is on the reporter plasmid under the control of a constitutive promoter (J23108) from the Anderson promoter library^60^ (plasmid map in Fig. S5b). Plasmids were constructed using the Golden Gate^61^ and Gibson assembly^62^ methods. Primers are listed in Table S2. Original OptoCre-VVD plasmids are from Sheets et al.^3^, where we use the OptoCre-VVD2 variant (Addgene #160400). The original FHY1 and ΔPhyA plasmids were constructed using the red light domains developed in Zhou et al.^41^. The *ho1-pcy* gene was derived from the pNO286-3 plasmid (Addgene #107746) that was deposited by Ong et al.^63^. We express Cre split with a photoreceptor pair as an operon (Fig. 1). The N-terminal fragment of Cre (nCre), amino acid AA(1-43), is followed by a 10 AA glycine-serine linker and FHY1 photoreceptor. A separate RBS is used to express ΔPhyA with a 10 AA glycine-serine linker to the C-terminal fragment of Cre (cCre). For experiments with alternative split sites, we split Cre with either nCre(1-136) or nCre(1-270).

### Light Exposure Assays and Recombinase Efficiency Characterization

In all experiments working with liquid cultures, cultures were maintained at 37 °C with continuous shaking at 220 rpm. On day one, an overnight culture was inoculated from a single colony and was incubated in LB medium supplemented with 100 μg/mL carbenicillin and 30 μg/mL kanamycin for plasmid maintenance. All cultures were shielded from surrounding light, except during designated red, blue, or ambient light exposure periods. The following day, cultures were diluted 1:100 in selective LB medium and induced with 100 μM IPTG, unless otherwise stated, for 2 hours prior to exposure to light. Light exposure was conducted using an OptoPlate-96 device equipped with LEDs emitting light at a wavelength of 630 nm (red) or 465 nm (blue) with an intensity of 155 μW/cm^2^ per well, unless otherwise stated. The OptoPlate-96 was constructed following the design by Bugaj et al.^64^ using the protocol by Dunlop^65^. Optical power measurements performed for different wells at different light intensity and wavelengths, and across multiple days demonstrated that variation in readouts of the intensity is modest (Fig. S6).

For light exposure conditions, cultures were subjected to red light exposure for a period of 4 hours unless stated otherwise. Following exposure, samples were transferred to selective LB medium devoid of IPTG for overnight growth. The following day, the fluorescence levels were measured using a BioTek Synergy H1m plate reader. For optical density (OD) readings in cells containing RFP, absorbance was conducted at 700 nm to avoid overlap with the RFP spectra, and the fluorescence read was conducted with an excitation of 584 nm and emission of 610 nm.

In experiments testing the impact of light intensity, cultures were exposed to red light for a total of 4 hours. Ambient light exposure involved exposing samples to laboratory lighting in a shaking incubator for either 5 minutes or 2 hours after IPTG induction, followed by continued incubation in the dark for the remainder of the 4 hours. Measurements of the ambient light in the laboratory had an intensity of 3.6 μW/cm^2^ at 630 nm.

In experiments involving blue light exposure, the same device and protocol were used. After 2 hours of IPTG induction, samples were exposed to blue light for a period of 4 hours. Then they were diluted 1:100 in selective LB media for overnight growth.

In experiments involving pulsatile red or blue light illumination, the OptoPlate-96 device was programmed to turn on red or blue LEDs for only 2-120 seconds every 5 minutes in an overnight experiment for a total period of 18 hours. The next day, all cultures were diluted 1:100 in selective LB medium and were incubated in the dark overnight. The following day, the fluorescence levels were measured using a BioTek Synergy H1m plate reader.

We confirmed target DNA excision genetically using PCR analysis and gel electrophoresis. We used a forward primer about 200 base pairs upstream of the first loxP site and a reverse primer about 100 base pairs downstream of the second loxP site to check for differences in band length before and after recombination. The primers are listed in Table S2.

### Microscopy and Image Analysis

We conducted single cell microscopy using a 100x objective and a Nikon Ti2-E microscope to analyze the bacteria containing OptoCre-REDMAP. After light stimulation, samples were diluted 1:100 for overnight growth in selective LB medium to allow full RFP expression. The next day, samples were diluted 1:100 2 hours prior to imaging in minimal media consisting of M9 salts supplemented with 2 mM MgSO4, 0.2% glycerol, and 0.01% casamino acids. After 2 hours, samples were placed on 1.5% low melting agarose pads made with the M9 minimal medium. Images were segmented and analyzed using DeLTA 2.0^66^.

### Minimum Inhibitory Concentration Measurement

Minimum inhibitory concentrations (MICs) were measured based on the protocol outlined in Sheets et al.^53^. Chloramphenicol antibiotic stocks were made by dissolving the antibiotic in 99% ethanol, with concentrations normalized for potency based on CLSI standards^67^. Assay plates for measuring the MIC were prepared by performing serial dilutions of antibiotic in 100 μL M9 minimal media (M9 salts supplemented with 2 mM MgSO4, 0.2% glycerol, 0.01% casamino acids, 0.15 μg/mL biotin, and 1.5 μM thiamine) in 96-well plates. Antibiotic concentrations were selected to include values that spanned the MIC levels for the dark and light state cultures in each experiment. Immediately following light exposure, cultures were diluted 1:100 and were transferred into 96-well plates in triplicate and grown overnight at 37°C. The OD absorbance reading of each well was then measured at 600 nm (OD600) using a BioTek Synergy H1 plate reader.

## Supporting Information

Supplementary figures with data supporting the manuscript. Supplementary tables with phantom optical properties and primer sequences.

## Acknowledgments

We thank Cristina Tous and Wilson Wong for providing us with the plasmids containing FHY1 and ΔPhyA, and Michael Sheets for helpful discussions on experimental design and light exposure protocols. We also thank Darren Roblyer and Carlos A. Gómez for providing the skin-mimicking phantom used in the light penetration experiments. This work was supported by NSF Grants MCB-2032357 and MCB-2324909 to MJD.

## Author Contributions

FJ and MJD conceived and designed the experiments. FJ performed the experiments and analyzed the data. FJ and MJD wrote the manuscript.

