## Supporting Information for "Red light responsive Cre recombinase for bacterial optogenetics"

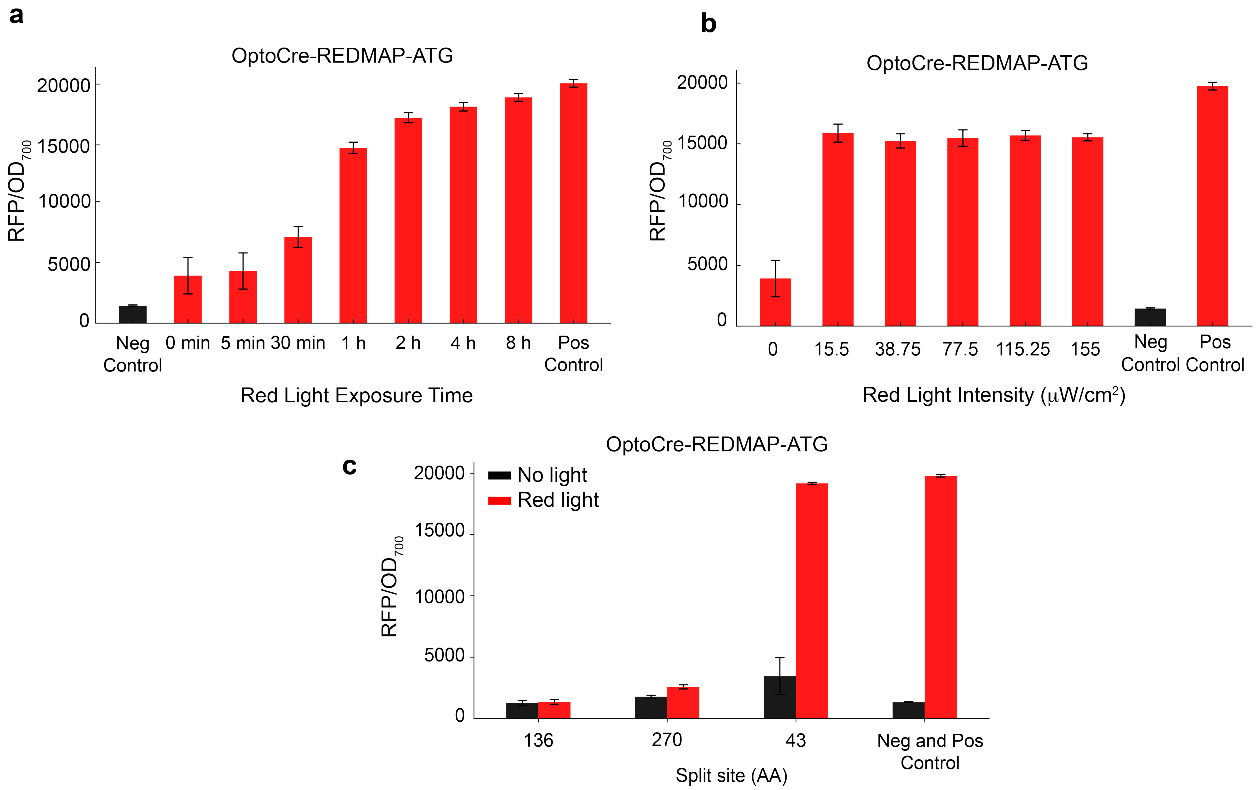


**Figure S1**. Characterizing the behavior of OptoCre-REDMAP (ATG version) and its split site variants. (a) Fluorescence measurements of RFP after different durations of red light exposure with OptoCre-REDMAP-ATG. (b) RFP output of OptoCre-REDMAP-ATG under red light intensities ranging from 15.5 to 155 μW/cm$^2$ with 4 hours of light exposure. (c) RFP output of split sites tested for OptoCre-REDMAP-ATG. AA, amino acid. Experiments use 4 hours of light exposure at 155 μW/cm$^2$. For all experiments in this figure, 100 μM IPTG induction was used. In all cases, error bars show standard deviation around the mean (n=3 biological replicates).

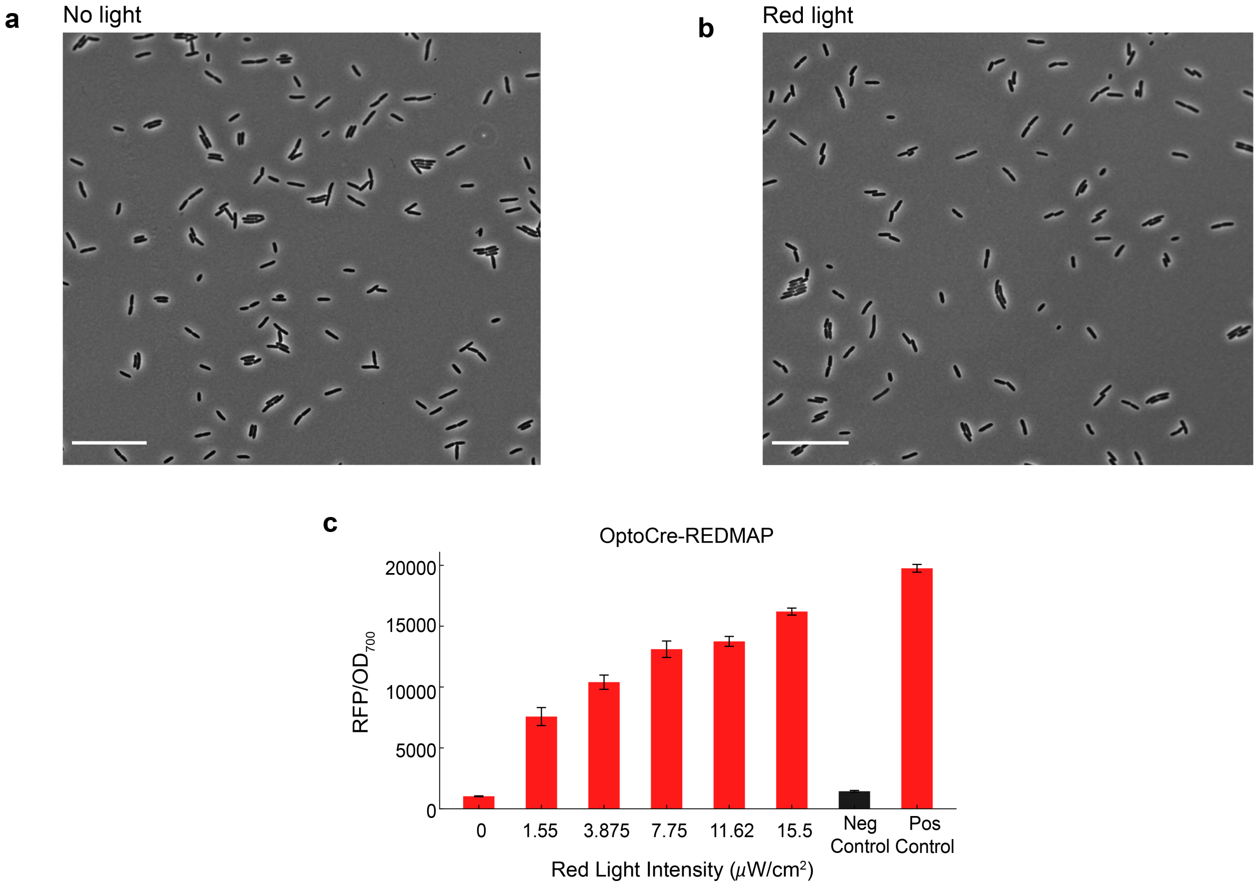


**Figure S2**. (a) Single cell phase contrast microscopy images of fully induced (100 µm IPTG) OptoCre-REDMAP cells without red light exposure, (b) and after 8 hours of 155 µW/cm$^2$ red light exposure. Scale bar = 10 µm. (c) RFP output in response to red light intensities ranging from 0 to 15.5 µW/cm$^2$ with 4 hours of light exposure. Error bars show standard deviation around the mean (n=3 biological replicates).

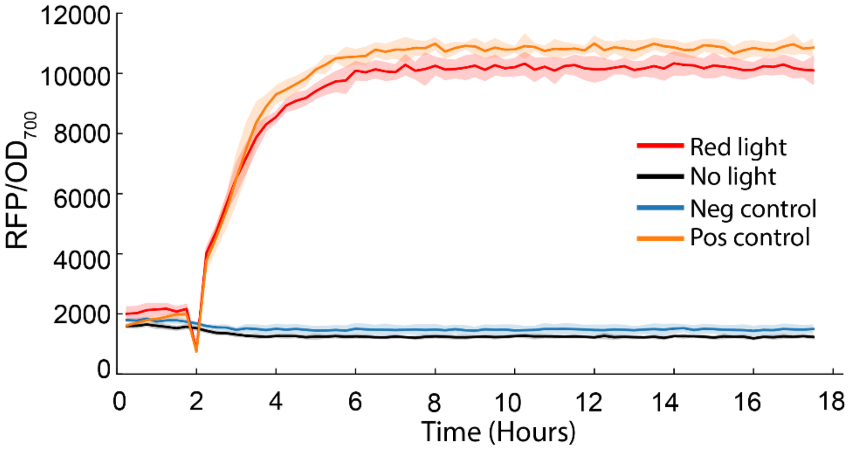


**Figure S3.** Real-time RFP expression of OptoCre-REDMAP in the dark or after being exposed to red light for 4 hours (t=-4 until t=0 hours), compared to the negative and positive control. Shaded error bars represent standard deviation around the mean from plate reader data (n=3 biological replicates).

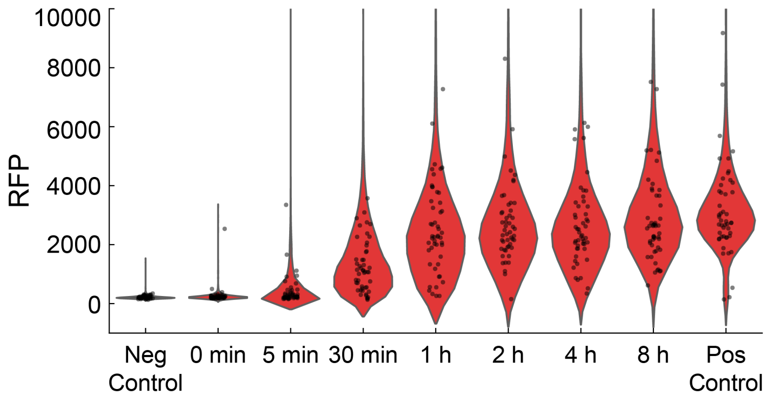


**Figure S4**. RFP distribution of single cells subjected to different red light exposure times ranging from 5 minutes to 8 hours, in addition to cells from negative control and positive control strains. Individual cell data are shown as dots on top of the violin plots.

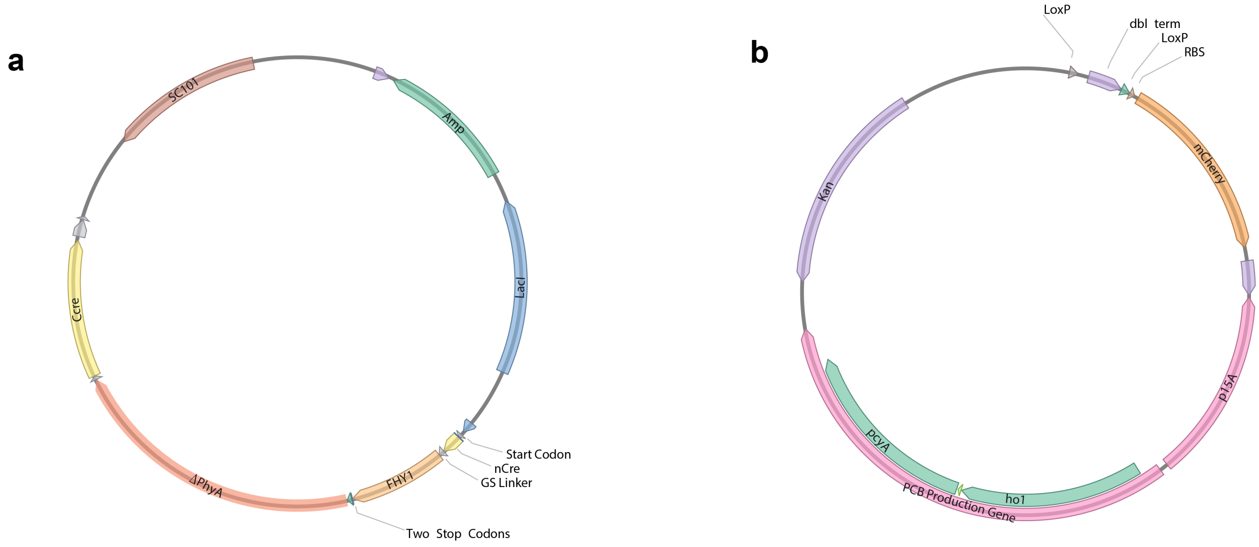


**Figure S5**. Plasmids maps. (a) OptoCre-REDMAP with the CTG start codon. (b) The reporter plasmid with a loxP flanked double terminator upstream of the gene encoding for mCherry. This plasmid also contains the *ho1-pcyA* gene encoding PCB.

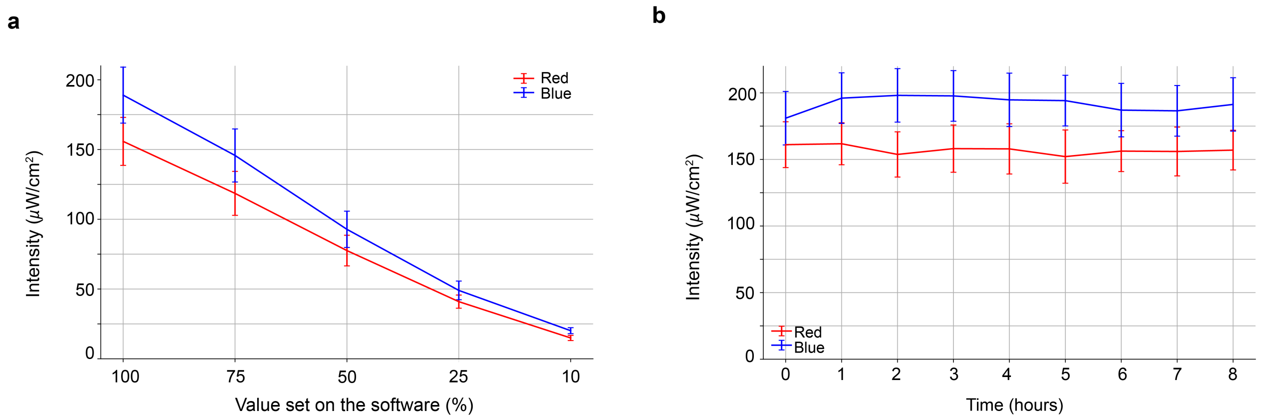


**Figure S6**. Optical power measurements of the OptoWell-96. (a) Light intensity values corresponding to red and blue LEDs measured at different intensities. (b) Light intensity values corresponding to red and blue LEDs when set to 100% intensity measured over an 8-hour period with 1 hour intervals. Error bars show standard deviation around the mean (n=3 replicates).

**Table S1**. Optical properties of silicone skin-mimicking phantom. $\mu_a$: absorption coefficient, $\mu_s'$: reduced scattering coefficient.

| **Phantom** | **Thickness (mm)** | **$\mu_a$ ($mm^{-1}$)** | | **$\mu'_s$ ($mm^{-1}$)** | |
|---|---|---|---|---|---|
| | | **730 nm** | **830 nm** | **730 nm** | **830 nm** |
| Phantom Skin | 1 | 0.010 | 0.007 | 1.62 | 1.27 |

**Table S2**. Primers used for cloning or sequencing.

| | |
|---|---|
| CCCGATCTTCCCCATCGG | PCR and sequencing primer to check terminator excision - Forward |
| CTCGAACTCGTGACCGTTAACAG | PCR and sequencing primer to check terminator excision – Reverse |
| AGGTAGGGTCTCGATCTGACAGCTAGCTCAGTC | Golden Gate PCR primer to amplify *ho1-pcy* gene - Forward |
| AGGTAGGGTCTCGCTATATAAACGCAGAAAGGCCC | Golden Gate PCR primer to amplify *ho1-pcy* gene - Reverse |
| AGGTAGGGTCTCCATAGCACCTGAAGTCAGCC | Golden Gate PCR primer to amplify reporter plasmid backbone for *ho1-pcy* insertion - Forward |
| AGGTAGGGTCTCCAGATTGCACTGAAATCTAGAAATATTTT | Golden Gate PCR primer to amplify reporter plasmid backbone for *ho1-pcy* insertion - Reverse |
| TTTCGTTTTCAGAGCAAGAG | PCR and sequencing primer to check the *ho1-pcy* gene insertion into the reporter plasmid - Forward |
| TTCCTCGTGCTTTACGGTAT | PCR and sequencing primer to check the *ho1-pcy* gene insertion into the reporter plasmid - Reverse |
| TCGGAGGTCTCCCAAGTGGTGGCAGCGGTACC CCCGAGGTGGAGGTCGAC | Golden Gate PCR primer to amplify *FHY1* gene - Forward |
| TCGGAGGTCTCCCCTCTTTAAAGTTAACTATTA CAGCATTAAAGTGCTAAAGTACTGCTC | Golden Gate PCR primer to amplify *FHY1* gene - Reverse |

| | |
|---|---|
| TCGGAGGTCTCGGAGGAGAAAGGTACCG CATGGAGAAGAAGATGAGCGGA | Golden Gate PCR primer to amplify Δ*phyA* gene - Forward |
| TCGGAGGTCTCGGAACCTCCGGTACC TTGGATTCCATCAATCTT | Golden Gate PCR primer to amplify Δ*phyA* gene - Reverse |
| TCGGAGGTCTCCGTTCAGGAGGTT CAGACAGATGCCAGGACATC | Golden Gate PCR primer to amplify the backbone plasmid containing Cre - Forward |
| TCGGAGGTCTCCCTTGATCCGGAGCCA GACCCAGAGTTCTCCATCAGGGA | Golden Gate PCR primer to amplify the backbone plasmid containing Cre - Forward |
| GGAGATATACATCTGGCCACCTCTGATGA | Gibson assembly PCR primer to amplify nCre section for changing start codon to CTG - Forward |
| TCCGGAGCCAGACCCGTTCAGCTTGCACCAGG | Gibson assembly PCR primer to amplify nCre section for changing start codon to CTG - Reverse |
| TGGTGCAAGCTGAACGGGTCTGGCTCCGGA | Gibson assembly PCR primer to amplify the OptoCre-REDMAP plasmid backbone for the insertion of CTG-nCre for changing start codon - Forward |
| ATCAGAGGTGGCCAGATGTATATCTCCTTCTTAAAAGATCTTT | Gibson assembly PCR primer to amplify the OptoCre-REDMAP plasmid backbone for the insertion of CTG-nCre for changing start codon - Reverse |
